# High quality genome assembly of the brown hare (*Lepus europaeus*) with chromosome-level scaffolding

**DOI:** 10.1101/2023.08.29.555262

**Authors:** Craig Michell, Joanna Collins, Pia K. Laine, Zsófia Fekete, Riikka Tapanainen, Jonathan M. D. Wood, Steffi Goffart, Jaakko L. O. Pohjoismäki

## Abstract

We present here a high-quality genome assembly of the brown hare (*Lepus europaeus* Pallas), based on a fibroblast cell line of a male specimen from Liperi, Eastern Finland. This brown hare genome represents the first Finnish contribution to the European Reference Genome Atlas pilot effort to generate reference genomes for European biodiversity.

The genome was assembled using 25X PacBio HiFi sequencing data and scaffolded utilizing a Hi-C chromosome structure capture approach. After manual curation, the assembled genome length was 2,930,972,003 bp with N50 scaffold of 125.8 Mb. 93.16% of the assembly could be assigned to 25 identified chromosomes (23 autosomes plus X and Y), matching the published karyotype. The chromosomes were numbered according to size. The genome has a high degree of completeness based on the BUSCO score (mammalia_odb10 database), Complete: 96.1% [Single copy: 93.1%, Duplicated: 3.0%], Fragmented 0.8%, and Missing 2.9%. The mitochondrial genome of the cell line was sequenced and assembled separately. The final annotated genome has 30,833 genes of which 21,467 code for a polypeptide.

The brown hare genome is particularly interesting as this species readily hybridizes with the mountain hare (*Lepus timidus* L.) at the species contact zone in northern Eurasia, producing fertile offspring and resulting in gene flow between the two species. In addition to providing a useful comparison for population studies, the genome can offer insight into the chromosomal evolution among Glires in general and Lagomorpha in particular. The chromosomal assembly of the genome also demonstrates that the cell line has not acquired karyotypic changes during culture.

## Introduction

The brown hare (*Lepus europaeus* Pallas), also known as the European hare, is a widespread species in the western parts of Eurasia (Bock, 2020). Besides its native range, the brown hare has been introduced to numerous regions, including the British Isles, the Falkland Islands, Canada, South America, Australia and New Zealand (Jaksic, 2023; Petrovan, 2013). In many places the brown hare is regarded as an invasive species and a threat to the local ecosystems (Stott, 2003) or native species (Reid, 2011).

The brown hare is a steppe-adapted species living in open grasslands and avoiding forested regions. Its colonization history in Europe is complex. The species has had glacial refugia during the Pleistocene in the Italian peninsula, the Balkans and Asia Minor (Fickel et al., 2008), with an interesting pre-glacial diversity hotspot in the archipelago of Greece (Minoudi et al., 2018). After the ice age, the species has been expanding its range both naturally as well as facilitated by human-caused changes in the landscape, especially through the expansion of agricultural lands and pastures. As mentioned above, the brown hare has been also frequently introduced by humans to new areas from antiquity to the present day (Petrovan, 2013), resulting in potential mixing of ancestral populations. This is true also for Finland, where the species has arrived naturally from the southeast through the Karelian isthmus in the late 19^th^ century (Ognev, 1940; Siivonen, 1972; Wadén, 1910) but also through local introductions to the southwestern parts of the country in the 1910s (Rikala, 1925; Salonius, 1924; Suomalainen, 1922; Wadén, 1910).

Curiously, Linné was unaware of the existence of the brown hare, despite having described the mountain hare (*Lepus timidus* L.) and the South African Cape hare (*Lepus capensis* L.). In fact, the brown hare is not native to Linné ‘s home country Sweden but was introduced there only a century after his death (Thulin, 2003). Curiously, the species author Peter Simon Pallas never published a formal description of the brown hare. Instead, the species was authority has been attributed to him because of inclusion of the name *Lepus europaeus* into his identification table of hare species of the world (Pallas, 1778: 30) (Figure 1A). Consequently, there is also no information of the type locality or any other information of the species in the original publication. The type locality of the brown hare has been retrospectively assigned – without too convincing arguments – to Burgundy (France) or Poland (Holden, 2005). Identifying Poland as the type locality was done by the Russian zoologist Sergey Ognev, and is based on Pallas mentioning in the 1778 text the hybrid forms of *Lepus variabilis* (synonym of *L. timidus*) and *L. europaeus* from Poland and Lithuania, as well as citing later works of Pallas for the species distribution (Ognev, 1940). In fact, Ognev (1940: 141) boldly states that “we consider southwestern Poland as a typical terrain for the nominal subspecies; there are hares with the characteristics of the basic form, expressed with full clarity” (translation from Russian).

**Figure 1.**
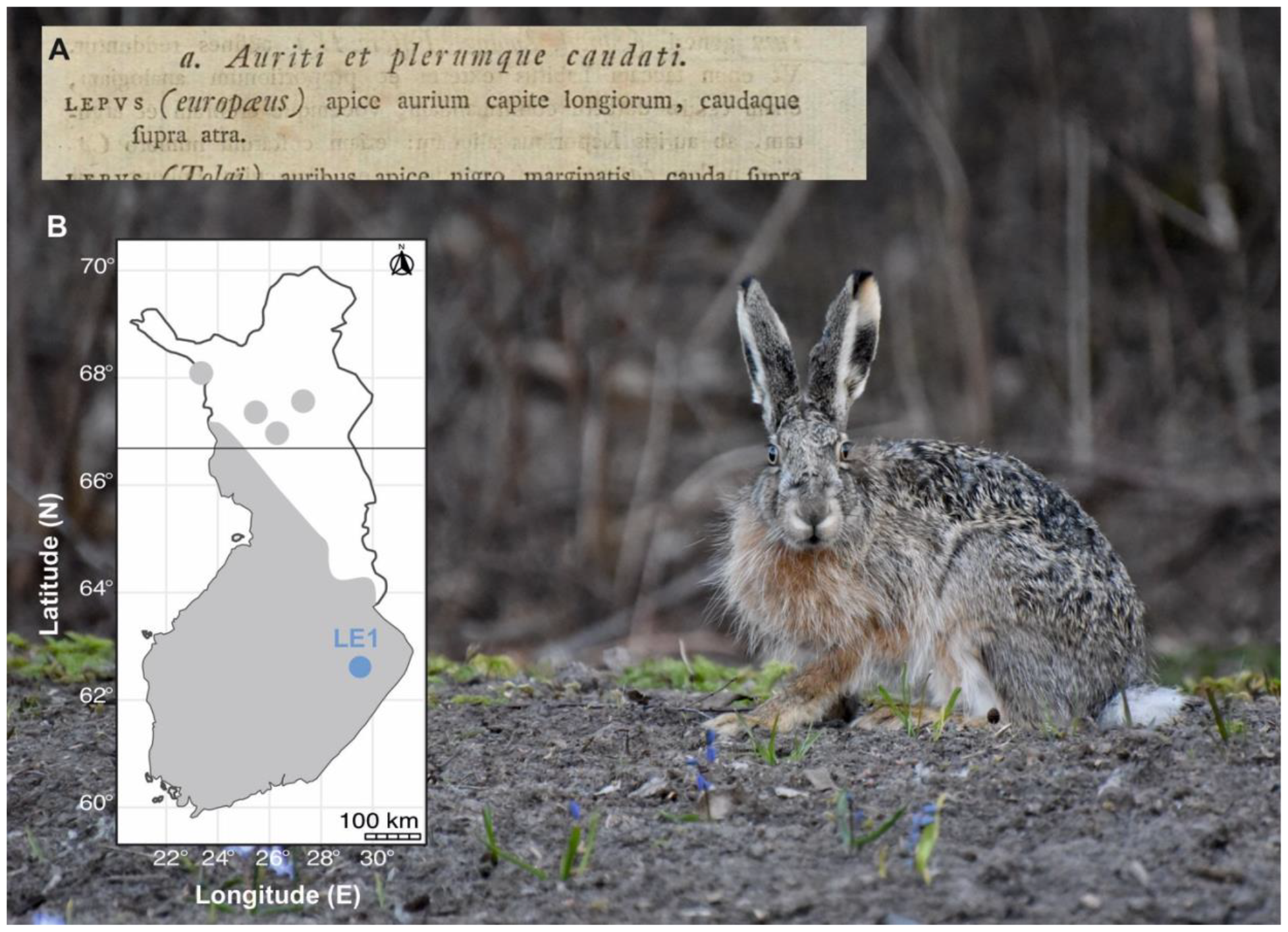
The brown hare. (A) An excerpt from the identification table by Pallas (1778: 30), which has been assigned as the species description for *Lepus europaeus*. “Tips of the ears longer than the head, tail black above.” (translation from Latin). (B) The geographic location of the LE1 sample used in this study. The grey-shaded area shows the approximate current distribution of the brown hare in Finland, based on the data in the Finnish Biodiversity Information Facility (FinBIF, https://laji.fi/en). Individual records from the north are from settlemens or towns. The arctic circle (black line) runs along the 66.6 ° parallel. Background photo shows a typical Finnish male brown hare. Photo taken in Joensuu, 20 km East from the LE1 sampling site by Dr. Mervi Kunnasranta.

While being widespread in the temperate regions of Western and Central Europe to the Caspian steppes in historical times, the brown hare began a northward expansion in the 19^th^ century. In the Fennoscandian region, the species reached the St Petersburg area by 1820s (Ognev, 1940), arriving in Finland at the turn of the century (Salonius, 1924; Siivonen, 1972; Thulin, 2003; Wadén, 1910). By 1930, the species had become established throughout southern Finland, with the range expansion stagnating southwest of the N64°-N62° -line until the 1990s. Quite likely benefitting from the ongoing anthropogenic climate change, brown hares range expansion has intensified during the last three decades, currently reaching the polar circle (Levänen et al., 2018a) (Figure 1B), as well as higher altitudes in the Alps (Schai-Braun et al., 2023). In Finland, the expansion seems to be limited with 150 days of annual snow cover (Levänen et al., 2018a). The range expansion together with the past introductions have brought the brown hare increasingly in contact with the mountain hare, especially at the northern edges of the species distribution, constituting a threat especially to the more temperate climate adapted populations and subspecies of the mountain hare (Reid, 2011; Reid and Montgomery, 2007; Thulin, 2003; Thulin et al., 2021). Besides competing, brown hares also hybridize with mountain hares, producing fertile offspring and resulting in gene flow between the species. This gene flow is biased towards the brown hare (Ferreira et al., 2021; Levänen et al., 2018a; Levänen et al., 2018b; Thulin and Tegelström, 2002), which obtains genetic variation from mountain hares, some of which might have adaptive significance (Pohjoismäki et al., 2021). In contrast to the brown hare ‘s expansion towards north, many of the Central European brown hare populations are contracting in range and numbers, likely driven by changes in agricultural practices and land use (Schai-Braun et al., 2013; Smith et al., 2005).

Overall, the brown hare ‘s features as an invasive species outside Europe and expansive species in the north, its hybridization tendency and complex population ancestries make the species highly interesting not only to study population genetics and adaptation mechanisms, but also for the understanding the genomic makeup of species boundaries (Gaertner et al., 2022). These studies would be greatly facilitated by the availability of high quality, chromosomally scaffolded reference genomes (Blaxter et al., 2022).

The last decade has witnessed a boom in genome sequencing technologies, leading to many sequencing initiatives, such as the Earth BioGenome Project (EBP), European Reference Genome Atlas (ERGA) and the Darwin Tree of Life (DToL), aiming to produce high quality reference genomes for a variety of organisms (Blaxter et al., 2022). This has been enabled by the technological development of high-throughput sequencing of long molecules. The read length N50 ‘s > 19 kbp, allows a high contiguity compared to the genome assemblies released in the early 2010 ‘s that were based on short read sequencing technologies. Additionally, Hi-C sequencing, a method measuring the contact frequency between all pairs of loci in the genome, has allowed producing pretext maps of chromosomes (Ghurye et al., 2019), enabling chromosome level assemblies with a precision that has been previously achieved only with model organisms (Lawniczak et al., 2022).

In the presented work, we apply the PacBio HiFi long-range sequencing technology with Hi-C sequencing to produce a high-quality reference genome of the brown hare (*Lepus europaeus*). While there is no previous genome assembly for the species, a mountain hare assembly (GCA_009760805) exists, representing the Irish subspecies *L. t. hibernicus* and is based on a female specimen, thus lacking the Y-chromosome. This assembly is also a so-called “pseudoreference “, assembled to chromosomes using the rabbit (*Oryctolagus cuniculus* L.) reference genome (Marques et al., 2020). The original rabbit reference genome had been established using whole-genome shotgun sequencing of females representing the inbred Thorbecke New Zealand White line and was quite fragmented (Carneiro et al., 2014), updated only recently with long-read sequencing data (Bai et al., 2021). Notably, the diploid chromosome number in domestic rabbits is 2n = 44 (Korstanje et al., 1999), whereas it is 48 in brown hare and mountain hare (Gustavsson, 1972).

To obtain a reference genome conforming to current standards, we used a fibroblast cell line (LE1) of a male specimen of brown hare (*Lepus europaeus*) from Liperi, Eastern Finland (Gaertner et al., 2022), as a source of high-molecular weight (HMW) DNA and fresh RNA. Compared to solid tissues, fibroblasts are optimal for Hi-C, as the method was originally developed for cells growing in a monolayer (Lieberman-Aiden et al., 2009). Now, over 240 years after the first mention of the name *Lepus europaeus* in the literature (Figure 1A), we have expanded the original seven-word description to encompass 2,930,972,003 genomic letters. These letters have been assembled in a highly complete (BUSCO mammalia_odb10 database, complete: 96.1%), and continuous (N50 scaffold: 125.8 Mb) manner with 93.16% of the sequence being placed in the identified 23 autosomes, and X and Y sex chromosomes. The presented genome assembly not only contains the identity of the brown hare as a species, but also provides a gateway to detailed knowledge of its biology and evolutionary history. Besides being the first Finnish contribution to the European Reference Genome Atlas (Mazzoni et al., 2023), the brown hare reference genome represents a continuing effort to map and understand our planets biodiversity.

## Methods

### Sampling and confirming of the species identity

A young male brown hare was sampled in October 2018 in Kuorinka, Liperi, Finland (62.6207 N, 29.4478 E, Figure 1B). In Finland, brown hares are highly dependent on similar man-made environments (Levänen et al., 2018a; Levänen et al., 2019), and also the collection location is agricultural area with a mosaic of fields, shrubs and mixed forest with a strong brown hare population and only occasional stray mountain hares. Geographically, the sampled population is part of the same distribution continuum as with species proposed type locality, through Russian Karelia and the Baltic states to Poland. This is important, as is recommendable that the species reference genome would represent or be closely related to the type locality population (Lawniczak et al., 2022).

The sampling had minimal impact on the local brown hare population and no impact on the habitat. As brown hares are legal game animals in Finland and the hunting followed the regional hunting seasons and legislation (Metsästyslaki [Hunting law] 1993/615/5§), the sampling adheres to the ARRIVE guidelines and no ethical assessment for the sampling was required. The sampling also did not involve International Trade in Endangered Species of Wild Fauna and Flora (CITES) or other export of specimens, as defined by the Convention on Biological Diversity (CBD). The species identity was confirmed at site based on the morphological features distinguishing the brown hare from the mountain hare, the only other hare species in northern Europe. Further analysis of the coding part of the LE1 genome and mitochondrial DNA haplotyping showed minimal ancestral admixture with mountain hares (Gaertner et al., 2022). In fact, unlike many other brown hares in the region, showing adaptive introgression of mountain hare specific UCP1 alleles (Pohjoismäki et al., 2021), the LE1 specimen is homozygous for the common ancestral brown hare allele, UCP02.

### Generation and vouchering of the cell line

The fibroblast cell line (LE1) was isolated from the specimen as described earlier (Gaertner et al., 2022) and is deposited as cryopreserved living cells under voucher number ZFMK-TIS-69747 into the biobank of the Stiftung Leibniz-Institut zur Analyse des Biodiversitätswandels, Zoological Research Museum Alexander König (ZFMK), Bonn, Germany. Additional identifiers for the sample are mLepEur2 and ERGA specimen ID ERGA_FI_3610_002 (COPO portal, https://copo-project.org).

### HMW DNA extraction and PacBio HiFi sequencing

A salting out-method was used to extract high molecular weight DNA from cells grown to confluency on a 10 cm cell culture dish. Briefly, the cells were detached from the dish using trypsin, followed by centrifugation and two washes with PBS. The salting out-method then followed the 10X Genomics “Salting Out Method for DNA extraction from cells” protocol (10x-Genomics, 2017). The high molecular weight DNA was quantified using a Qubit 3.0 (Thermo Fisher Scientific™, Malaysia), followed by qualification with an 0.8 % Agarose gel. The DNA was sent for the library preparation and sequencing at the DNA Sequencing and Genomics Laboratory, Institute of Biotechnology, University of Helsinki on the PacBio Sequel II. In brief, the DNA was quantified using Qubit and DNA fragment length was measured using Fragment Analyzer (HS large fragment kit). DNA was then sheared with Megaruptor 3 (Diagenode) (45/200/30) to produce DNA of length 24 kb. Buffer was then replaced with PacBio ‘s Elution buffer using AMPure beads. Repair and A-tailing, adapter ligation, nuclease treatment and cleanup with SMRTbell cleanup beads was then performed with the SMRTbell prep kit 3.0. Larger fragments (greater than 10 000 bp) were then purified with the BluePippin (Sage Science) (0.75% DF Marker S1 high pass 6-10kb v3) and DNA was further purified with PacBio AMPure beads and treated with DNA damage repair mix (PacBio) (37°C for 60 mins) and then purified with AMPure clean-up beads and eluted to 11 ul PacBio ‘s Elution buffer. Libraries were sequenced on the PacBio Sequel II at a concentration of 90 pM using PacBio ‘s instructions (provided by the SMRTlink software). Sequence data from two flow cells was produced. Genomescope2 was used to visualize the HiFi sequencing results (Ranallo-Benavidez et al., 2020).

### Mitochondrial DNA (mtDNA) sequencing

The mitochondrial genomes of our hare cell lines (Gaertner et al., 2022) have been sequenced for the purposes of another study using approximately 2 kb overlapping PCR-amplified fragments of mtDNA. The primers used to amplify the mountain hare mtDNA were as follows:

Le93F: TTGTTTTGTAGCAAGTTTACACATGC

Le184R: GCTTAATACCTGCTCCTCTTGATCTA

Le1580F: TTAAACCCATAGTTGGCCTAAAAGC

Le1635R: TTGAGCTTTAACGCTTTCTTAATTGA

Le3045F: AGGCGTATTATTTATCCTAGCAACCT

Le3175R: CCTCATAAGAAATGGTCTGTGCGA

Le3921F: CCCCCTAATCTTTTCCATCATCCTAT

Le4482R: TCATCCTATATGGGCAATTGAGGAAT

Le4689F:AGGCTTTATTCCAAAGTGAATTATTATTCA

Le5417R: AGGCTCCAAATAAAAGGTAGAGAGTT

Le6696F: ATACCGTCTCATCAATAGGCTCCTTC

Le6756R:ATAAAGATTATTACTATTACAGCGGTTAGA

Le8603F: AGCCTATATCTACATGATAATACTTAATGA

Le8698R: CGGATAAGGCCCCGGTAAGTGG

Le10552F: TTGAAGCAACACTAATCCCTACACTA

Le10613R: TCGTTCTGTTTGATTACCTCATCGT

Le11301F: ACCATTAACCTTCTAGGAGAGCTTCT

Le11807R: AGGATAATGATTGAGACGGCTATTGA

Le12407F: GTCTAATCCTAGCTGCTACAGGTAAG

Le12791R: GAGCATAAAAAGAGTATAGCTTTGAA

Le14204F: ATTGTTAACCACTCTCTAATCGACCT

Le14514R: CCAATGTTTCAGGTTTCTAGGTAAGT

Lt16056F: TGGGGTATGCTTGGACTCAAC

Le16119R: TCGTCTACAATAAGTGCACCGG

In total, 12 separate reactions were prepared to cover the mitochondria genome:

1. Lt16056F + Le184R: 1871 bp
2. Le93F + Le1635R: 1543 bp
3. Le1580F + Le3175R: 1596 bp
4. Le3045F + Le4482R: 1438 bp
5. Le3921F + Le5417R: 1497 bp
6. Le4689F + Le6756R: 2068 bp
7. Le6696F + Le8698R: 2003 bp
8. Le8603F + Le10613R: 2011 bp
9. Le10552F + Le11807R: 1256 bp
10. Le11301F + Le12791R: 1491 bp
11. Le12407F + Le14514R: 2108 bp
12. Le14204F + Le16119R: 1916 bp

(Expected fragment size based on the published *Lepus europaeus* mtDNA sequence from Sweden [NC_004028.1]).

The fragments were amplified from total DNA preparations using a PCR program with a 1 min 94 °C denaturing step, followed by 35 cycles of 94 °C for 15 s, 56 °C for 15 s and 72 °C for 2 min and a final 3 min elongation step at 72 °C. The obtained products were gel purified using the GeneJET gel extraction kit (Thermo Fisher Scientific™) and sent for sequencing using Illumina MiniSeq at the Genome Center of Eastern Finland.

The sequence of the non-coding region-containing PCR fragment (Lt16056F + Lt184R) was further validated by Sanger sequencing, applying also the following additional sequencing primer:

Le101F: TATAAATTCCTGCCAAACCCCAAAAA

### Hi-C library preparation

Hi-C sequencing libraries were prepared following the protocol of (Belaghzal et al., 2017) with the following changes: 1.) To prepare a diverse sequencing library, we performed the Hi-C protocol in triplicates, 2.) Size fractionation was performed using the NEBNext Ultra II FS DNA module (New England Biolabs), 3.) Library preparation was performed using the NEBNext Ultra II library preparation kit for Illumina (New England Biolabs) and 4.) Library enrichment was performed using triplicate PCR reactions with six cycles of PCR. The PCR reactions were then purified using Ampure XP beads (Beckman Coulter Life Sciences) at a ratio of 0.9X. The final clean libraries were quantified using Qbit, followed by agarose gel electrophoresis to confirm the fragment size. The sequencing was performed on a single lane of the Illumina NovaSeq 6000 using the SP flowcell with paired-end chemistry 2 x 150bp.

### Genome assembly

HiFiasm version 0.18.7 (Cheng et al., 2021) was used to assemble the 25X (rq >0.99) PacBio HiFi reads using the recommended arguments *-l3 –h1 and –h2 –primary* to integrate the Hi-C read data and produce a primary assembly. We then continued with the scaffolding of the primary assembly and annotation. To process the HiC data, we first mapped the Hi-C data to the primary genome assembly using BWA-mem version 0.7.17 (Li and Durbin, 2009) with the arguments *-SP*. The mapped reads were then parsed and filtered using pairtools version 1.0.2 (Open2C et al., 2023). To parse the aligned Hi-C reads, we used the options *–no-flip –dd-columns mapq –walks-policy mask*. The parsed pair data was then sorted and deduplicated using default arguments with pairtools. Finally, we selected unique-unique and unique-rescued pairs and split these into the pairs file and bam file for input in YaHS version 1.1 (Zhou et al., 2023). YAHS was run using the default parameters with the primary contig assembly and the filtered Hi-C bam file. HiFiasm operates as a haplotype-aware assembler, counting and assembling unique k-mers. Utilizing k-mer coverage information, the assembler identifies regions in the genome with expected heterozygous coverage and subsequently amalgamates them into a unified sequence. To refine the assembly further, Hi-C data is incorporated to elucidate the phasing of variants and ascertain connected genomic regions. Consequently, the assembler ‘s output provides a primary assembly along with an alternative assembly, effectively separating the haplotypes by their distinct alleles within the genomic content (please see https://lh3.github.io/2021/04/17/concepts-in-phased-assemblies). Contiguity and general genome statistics were calculated using QUAST version 5.2.0 (Mikheenko et al., 2018). We assessed the completeness of the genome by calculating the number of complete single copy orthologs with BUSCO version 5.1.2 (Manni et al., 2021), using the mammalia_odb10 database as well as the more lineage specific glires_odb10 database.

### Genome annotation

Repeat annotation of the genome was performed with EDTA version 2.1.0 (Ou et al., 2019), a *de novo* repeat identification pipeline. Using the repeat library produced by EDTA, we masked the scaffolded genome using RepeatMasker version 4.1.1 (Smit et al., 2013-2015). RNA-seq data from the same cell line (SRA accession number: SRR18740842) as well as RNA-seq data from other *L*.*timidus* libraries (SRA accession number: SRR10020054, SRR10020055, SRR10020060, SRR10491719, SRR18740839, SRR18740840, and SRR18740841) was collected from the sequence read archive (SRA). The previously produced RNA-seq data (Gaertner et al., 2022) was trimmed using fastp version 0.23.2 (Chen et al., 2018) and mapped against the masked genome using HISAT2 version 2.1.0 (Kim et al., 2019) with the default parameters. Furthermore, we included the protein sequences of the assembled transcripts from this cell line as further evidence for the genome annotation. These lines of gene evidence were included in the annotation using BRAKER3 version 3.02 (Bruna et al., 2021). The original annotation has been further augmented by the NCBI Eukaryotic Genome Annotation Pipeline (https://www.ncbi.nlm.nih.gov/genome/annotation_euk/process/). Telomeric sequences [AACCCT]n were identified using a Telomere Identification toolkit (tidk) version 0.2.31. The telomeric sequence copy number was then calculated in windows of 200kb for visualization using Circos (Krzywinski et al., 2009).

### Mitochondrial DNA assembly and annotation

Mitochondrial DNA was assembled from the PCR-amplified and Illumina sequenced mtDNA using the MitoZ pipeline (Meng et al., 2019). After comparing the results of the pipeline ‘s outputs with different kmer options, we selected the best assembly as final. Run options used in the final assembly were --clade Chordata –fastq_read_length 150, --requiring_taxa Chordata -- genetic_code 2 --kmers_megahit 21 29 39 59 79 99 119 141. The tools invoked by the pipeline included fastp (Chen et al., 2018) for cleaning the raw data, MEGAHIT (Li et al., 2015) for assembly, after which sequences were filtered to ensure the correct taxa by HMMER (Wheeler and Eddy, 2013) and further filtered for completeness of protein-coding genes. Annotation steps were done using TBLASTN (Gertz et al., 2006), GeneWise (Birney et al., 2004) and MiTFi (Juhling et al., 2012). Final manual curation, the annotation of the non-coding region (NCR) as well as the illustration of the mitochondrial genome was done using Geneious® 10.2.6 (Biomatters. Available from https://www.geneious.com). The functional loci on the NCR were identified based on the similarity with the human (NC_012920) and mouse (FJ374652) NCR sequences.

### Manual curation

The assembled and annotated genome was manually curated to further improve its quality based on Hi-C contact maps as described in (Howe et al., 2021), allowing to break or join erroneus scaffold assemblies and removing duplicated contigs.

### Comparison with previous assembly

We performed a comparison of our scaffolded assembly to the current *L. timidus* genome assembly (NCBI Accession number: GCA_009760805). Mapping to the genome was performed using minimap2 version 2.21 (Li, 2018) with the arguments *-asm5*. A dot plot of the alignment was created using the R script pafCoordsDotPlotly.R (https://github.com/tpoorten/dotPlotly).

## Results

The genome assemblies can be accessed via BioSample accession SAMN33984520 as well as BioProject accession PRJNA1009711 for the primary assembly and PRJNA1009710 for the alternative assembly. The Final annotated assembly can be found under NCBI Reference Sequence assembly accession GCF_033115175.1, which can be viewed also using the NCBI Genome Data Viewer (https://www.ncbi.nlm.nih.gov/genome/gdv/browser/genome/?id=GCF_033115175.1).

### Genome assembly

The expected haploid genome size of the *L. europaeus* is similar to *L. timidus*, which has been estimated to be 3.25 pg (3,178,500,000 bp) (Vinogradov, 1998), containing 23 autosomal chromosomes and two sex chromosomes (Gustavsson, 1972). PacBio HiFi sequencing with two flow cells resulted in 25 X coverage of the expected genome size. The sequence N50 of the HiFi data was 19.97 kb. When k-mer size of 21 is used, the resulting genome size from the PacBio HiFi data is 2.4 Gbp (Figure 2A). The Illumina sequencing of the Hi-C data produced 494,285,585 paired reads, representing about 51 X coverage of the genome, with a duplication rate of 17 %. Assembly with HiFiasm yielded a contig assembly of 2.96 Gbp made up of 671 contigs with a contig N50 of 58 Mbp. The longest contig was 149 Mbp. Using the uniquely mapped Hi-C data, we were able to scaffold the contigs and fix misassembled contigs (Figure 2B). Although the genome size (2.96 Gbp) and the largest scaffold (149 Mbp) were similar to the contig assembly, the scaffold N50 was increased to 124 Mbp (N=10) after the Hi-C scaffolding. The obtained scaffold N90 was 21 Mbp (N=25). The BUSCO scores of the *L. europaeus* assembly suggest it to be near-complete, with the following results (%-of the total genome): Complete: 96.1% [Single copy: 93.1 %, Duplicated: 3.0 %], Fragmented 0.8 %, and Missing 2.9 % based on the mammalia_odb10 database.

**Figure 2.**
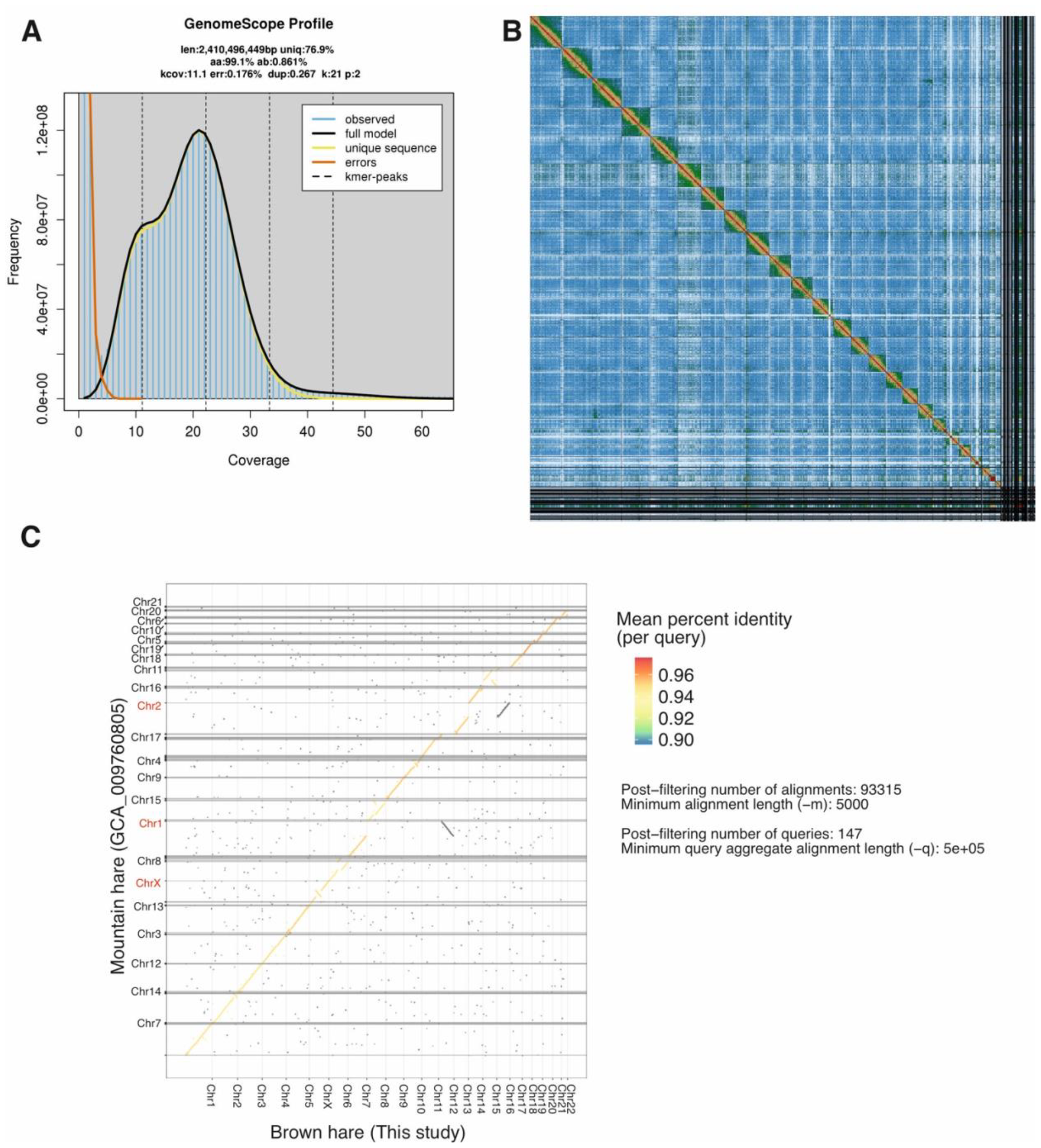
A) Genomescope2 profile of the PacBio HiFi data. B) YaHS pretext map of Hi-C scaffolding after manual curation. C) Dot plot comparison with the mountain hare pseudoreference (GCA_009760805) assembly with rabbit chromosomal assignments. Each grey horizontal line represents a contig end. Names of the non-chromosomal contigs have been omitted for clarity. Rabbit chromosomes corresponding two hare chromosomes as well as the X chromosome with an inversion are are highlighted.

To further improve the assembly, manual curation was performed, resulting in 76 scaffold breaks, 81 joins and removal of 62 erroneously duplicated contigs (Table 1). These interventions led to an increase in scaffold N50 of 1.69 % from 123.7 Mb to 125.8 Mb, and a reduction in scaffold count of 9.21 % from 716 to 650. Of the finalized assembly, 93.16 % could be assigned to 25 identified chromosomes (23 autosomes plus X and Y), matching the known karyotype. Chromosomes were numbered according to size (Table 2). The curated genome length was 2,930,972,003 bp.

**Table 1.** The brown hare reference genome chromosome assignment and assembly statistics.

| **Chromosome assignment** |  |
| --- | --- |
| **Chr length** | 2,730,543,680 bp |
| **Autosomes** | 23 |
| X | 1 |
| Y | 1 |
| Unlocalised | 19 |

| **Statistics for primary assembly**<br>**after manual curation** |  |
| --- | --- |
| **# scaffolds** | 650 |
| Total scaffold length | 2,930,972,003 |
| **Average scaffold length** | 4,509,187.70 |
| **Scaffold N50** | 125,776,599 |
| **Scaffold auN** | 117,151,857.99 |
| **Scaffold L50** | 10 |
| Largest scaffold | 184,151,585 |
| **Smallest scaffold** | 1000 |
| **# contigs** | 1068 |
| Total contig length | 2,930,889,045 |
| **Average contig length** | 2,744,278.13 |
| **Contig N50** | 43,409,466 |
| **Contig auN** | 47,572,960.95 |
| **Contig L50** | 21 |
| Largest contig | 141,329,000 |
| **Smallest contig** | 1000 |
| **# gaps in scaffolds** | 418 |
| Total gap length in scaffolds | 82,958 |
| Average gap length in scaffolds | 198.46 |
| **Gap N50 in scaffolds** | 200 |
| Gap auN in scaffolds | 199.70 |
| **Gap L50 in scaffolds** | 208 |
| Largest gap in scaffolds | 200 |
| Smallest gap in scaffolds | 31 |
| Base composition |  |
| **A** | 821,587,687 |
| **C** | 645,897,412 |
| **G** | 644,604,632 |
| **T** | 818,799,314 |
| **GC content** | 44.03 % |
| **# soft-masked bases** | 0 |
| **# segments** | 1,068 |
| **Total segment length** | 2,930,889,045 |
| **Average segment length** | 2,744,278.13 |
| **# gaps** | 418 |
| **# paths** | 650 |

**Table 2.**
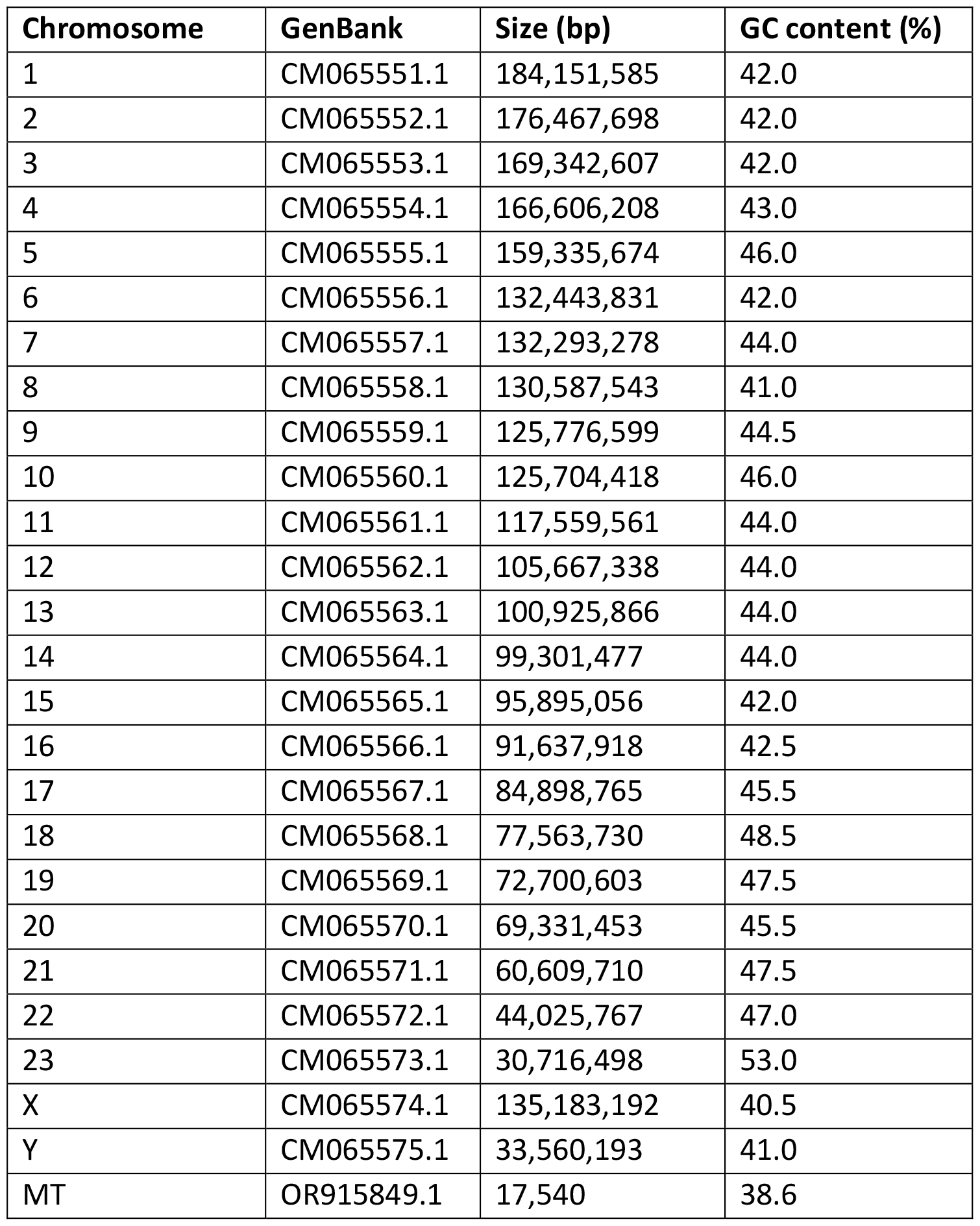
Chromosome assemblies of the brown hare reference genome with their corresponding GenBank access numbers.

| **Chromosome** | **GenBank** | **Size (bp)** | **GC content (%)** |
| --- | --- | --- | --- |
| 1 | CM065551.1 | 184,151,585 | 42.0 |
| 2 | CM065552.1 | 176,467,698 | 42.0 |
| 3 | CM065553.1 | 169,342,607 | 42.0 |
| 4 | CM065554.1 | 166,606,208 | 43.0 |
| 5 | CM065555.1 | 159,335,674 | 46.0 |
| 6 | CM065556.1 | 132,443,831 | 42.0 |
| 7 | CM065557.1 | 132,293,278 | 44.0 |
| 8 | CM065558.1 | 130,587,543 | 41.0 |
| 9 | CM065559.1 | 125,776,599 | 44.5 |
| 10 | CM065560.1 | 125,704,418 | 46.0 |
| 11 | CM065561.1 | 117,559,561 | 44.0 |
| 12 | CM065562.1 | 105,667,338 | 44.0 |
| 13 | CM065563.1 | 100,925,866 | 44.0 |
| 14 | CM065564.1 | 99,301,477 | 44.0 |
| 15 | CM065565.1 | 95,895,056 | 42.0 |
| 16 | CM065566.1 | 91,637,918 | 42.5 |
| 17 | CM065567.1 | 84,898,765 | 45.5 |
| 18 | CM065568.1 | 77,563,730 | 48.5 |
| 19 | CM065569.1 | 72,700,603 | 47.5 |
| 20 | CM065570.1 | 69,331,453 | 45.5 |
| 21 | CM065571.1 | 60,609,710 | 47.5 |
| 22 | CM065572.1 | 44,025,767 | 47.0 |
| 23 | CM065573.1 | 30,716,498 | 53.0 |
| X | CM065574.1 | 135,183,192 | 40.5 |
| Y | CM065575.1 | 33,560,193 | 41.0 |
| MT | OR915849.1 | 17,540 | 38.6 |

### Genome annotation

EDTA produced a curated custom repeat library of 7,045 repetitive elements. Interestingly, the proportion of the genome masked as repetitive elements was higher than expected at 46.8 % of the genome. In the mountain hare pseudoreference genome assembly (GCA_009760805), only 28 % of the genome was masked, and similarly the *k*-mer based estimate of the repetitive elements was 26.7 %. To annotate the genome, we mapped ∼1.6 billion RNA-seq reads from previously published RNA-seq libraries, including this cell line. The average mapping rate of all libraries was 89.95 %, which we consider a good indicator of the quality of this genome. Using BRAKER3, we were able to annotate 19,906 gene models in the genome. We then ran BUSCO on the predicted genes and the results show a good level of completeness: Complete: 81.1 % [Single copy: 63.3 %, Duplicate: 17.8 %], Fragmented: 1.1 %, Missing: 17.8 %, Total: 9226 when compared to the mammalia_odb10 database. The final annotation (NCBI RefSeq GCF_033115175.1-RS_2024_01) contains 30,833 genes of which 21,467 code for a polypeptide. Telomeric sequences on both ends of the chromosome were detected on 13 chromosomes (Figure 3A). Furthermore, telomeric sequences were found in high copy number throughout the chromosome length, a feature previously noted using FISH in mountain hares (Forsyth et al., 2005).

**Figure 3.**
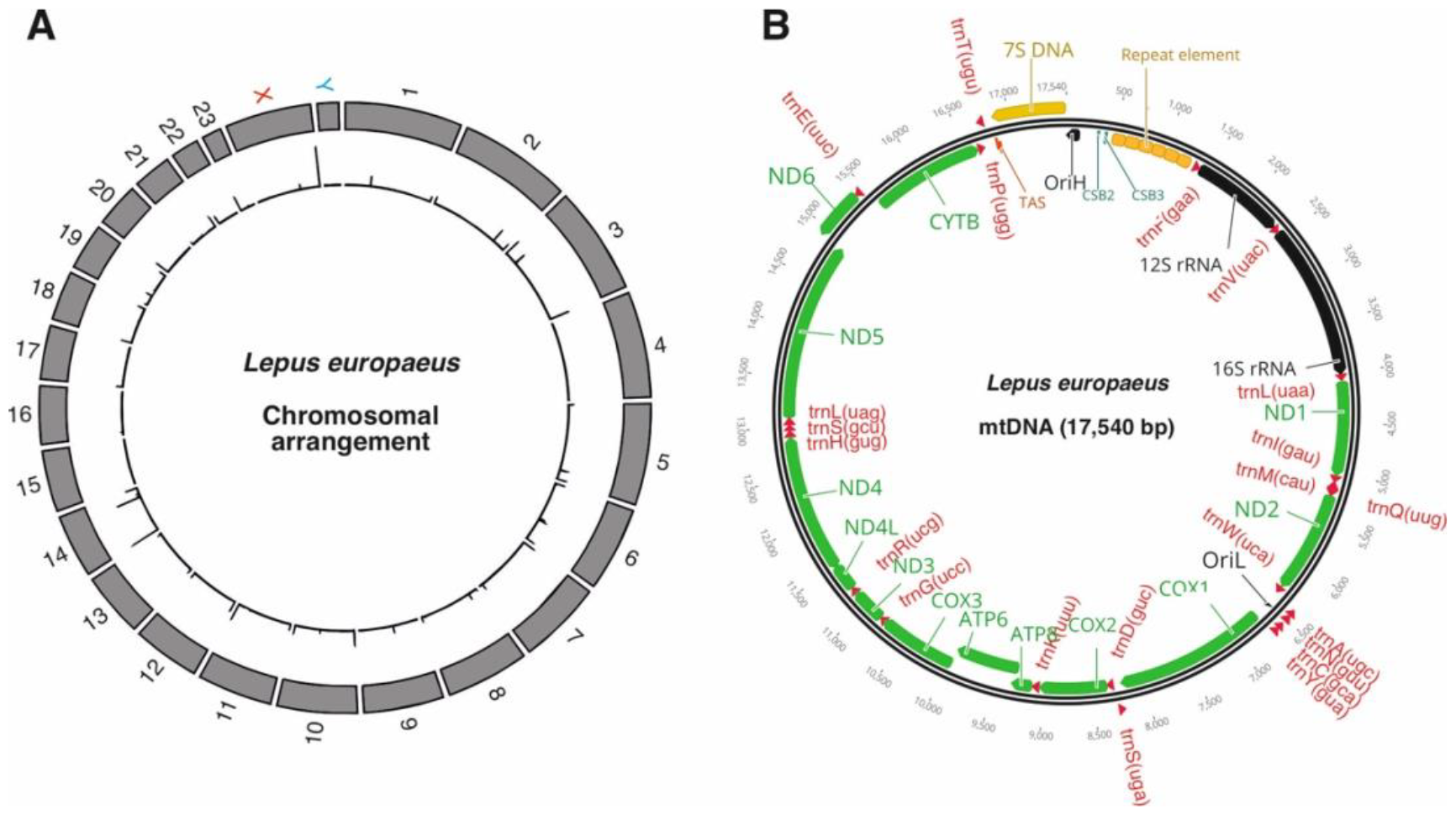
A reference genome for the brown hare. A) A size map of 23 autosomes as well as X and Y sex chromosomes of *Lepus europaeus*. Telomeric sequences and their relative lengths are indicated on the inner circle. B) A schematic illustration of the mountain hare mitochondrial genome. Genes encoded by the H-strand are placed inside and the L-strand genes outside of the circle. NCR = non-coding region.

### Comparison to the mountain hare assembly

Minimap2 was able to align 97.3 % of the contig sequences from the mountain hare pseudoreference genome assembly (GCA_009760805) to this genome assembly (Figure 2C). As pointed out earlier, the mountain hare genome assembly has been scaffolded against the rabbit reference genome (Giska et al., 2022), which has a different chromosome count (Beklemisheva et al., 2011) and explains the few notable inversions and structural differences between the two genomes (Figure 2C). For example, the chr7 and chr 12 assemblies of the brown hare have a good degree of synteny with the mountain hare pseudoreference (= rabbit) chr1, with the chr12 being in inverted orientation. Similarly, the brown hare chr13 and chr16 assemblies have a good synteny and orientation with the rabbit chr2.

### Mitochondrial genome

While it is possible to recover the entire mitochondrial genome from the HiFi sequencing data (Uliano-Silva et al., 2023), we had already previously sequenced the mitochondrial genomes of our hare cell lines for the purposes of another study. The assembly of the Illumina data provided a 16,836 bp mtDNA sequence, which was complete and circular, with a 142 bp non-repetitive overlap between the 3^’^ and 5^’^ ends of the original assembly removed from the final version. All the expected vertebrate mitochondrial genes, 37 in total, were found (Figure 3B).

Interestingly, the primary assembly of the Illumina data was notably smaller than expected based on the original PCR amplicons, with a notable difference between the expected and obtained non-coding region (NCR) length. Sanger sequencing of the PCR-fragment containing the NCR (Lt16056F + Le184R) revealed that it contained a sequence run consisting of six head-to-tail repeats of a 140 bp element between OriH and tRNA-Phe, explaining the incorrect assembly of the NCR sequence from the short-read Illumina data. With the addition of the repeat elements, the final mitochondrial genome size of our specimen is 17,540 bp.

Based on the sequence similarity with human and mouse mitochondrial genomes, we were able to tentatively identify the termination associated sequence (TAS, 2 nt difference to human), required for the generation of the 7S DNA and thus the displacement loop (D-loop) structure on the NCR (Crews et al., 1979). Interestingly, assuming that the 7S DNA is roughly the same size as in other mammals (600-650 nt), its 5 ‘-end would map to the beginning of another highly repetitive 240 bp region, likely to have structural significance. This is a notably longer sequence run than the tRNA-like sequence reported at the replication origin of *L. timidus* mtDNA (Melo-Ferreira et al., 2014). We have preliminary assigned this structure to correspond to the heavy-strand origin of replication (OriH), supported by the fact that a similar, tRNA sequence-derived hairpin structure is required for the priming of the origin of the light-strand replication (OriL) (Fuste et al., 2010). Experimental validation of the proposed OriH is required.

## Discussion

Given the challenges of assembling a large mammalian genome, we have produced a high-quality genome assembly for the brown hare. The genome assembly has a high degree of contiguity (Contig N50 43 Mbp) and completeness (BUSCO complete 96.1 %). Interestingly, the genome contains a high number of repetitive elements, a fact that warrants further investigation to elucidate their identity. While vast majority of the sequences seem to come from retroelements or endoretroviruses, we cannot fully exclude that some multicopy host genes could be masked. A detailed analysis of the repeat element libraries from this and another vertebrate genome assembly releases might be warranted.

The genome assembly is in chromosome scale, and all 23 autosomes as well as X and Y chromosomes could be scaffolded. Compared with the closely related mountain hare pseudoreference assembly, the brown hare genome assembly is much more contiguous (Table 1) and inclusion of telomeric repeats on both ends of several chromosomes (Figure 3A) further demonstrates the telomere-to-telomere continuity of these assemblies. The presented assembly also resolves the expected chromosome structure for the species (Figure 2B, C), which is informative regarding the karyotype difference between hares (n = 24) and rabbits (n = 22). Comparison of the brown hare assembly with the mountain hare pseudoreference chromosomes shows that the rabbit chromosome 1 corresponds to the brown hare chromosomes 12 and 17, with the chromosomal arm corresponding to the former is in an inversed orientation. Similarly, the rabbit chromosome 2 corresponds to the hare chromosomes 13 and 16, both in the same orientation. It should also be noted that as the hare chromosomes were numbered according to their size, the numbering in general does not match those of the rabbit chromosomes (Figure 2C)

The obtained 17,540 bp long mitochondrial genome assembly also provides some interesting aspects of mtDNA sequence variation within mammalian species. The size difference to human (16,569 bp) and mouse (16,298 bp) mitochondrial genomes is caused by a rather long non-coding region (2,102 bp vs. 1,123 bp and 879 bp in humans and mice, respectively), longer rRNA and tRNA genes, as well as additional non-coding nucleotides between genes. The mitochondrial genome of our specimen is slightly smaller than the 17,734 bp previous NCBI Reference Sequence for brown hare (NC_004028.1) from Sweden. The size difference is mostly due to a copy number difference in tandem repeat elements within the non-coding region, with seven in the NC_004028.1 and six in our specimen. Indels elsewhere in the NCR explain the rest of the difference.

The main technical challenge for highly continuous, good quality reference genome assemblies is the availability of relatively large quantities of intact HMW DNA for HiFi sequencing. This typically requires snap freezing of tissue samples and assuring an intact cold chain of below -80 °C for their sending, handling, and long-term storage (Januszczak and Holroyd, 2023). While development in sequencing technologies can reduce the required DNA amounts, obtaining high sequencing coverage over a large vertebrate genome is easier when material is plentiful. Similarly, RNA is very sensitive to degradation in post-mortem tissue samples, also requiring immediate preparation or snap-freezing to be suitable for RNA-seq. Although transcript data is not obligatory, it is highly useful for the purpose of genome annotation. The requirement of large samples of fresh tissue for -omics purposes complicates the sampling of most vertebrates species. For example, in Finland all land vertebrates are protected by law (animal protection law 247/1996, hunting law 615/1993), with very few exceptions, such as game animals and pest species. Capturing and lethal sampling is especially problematic for endangered species. Not only will the sampling require extensive permits, but it is also difficult to justify and conduct ethically. Here, the generation of primary fibroblast cell lines can offer a solution. Although the source specimen for the cell line utilized in this study was hunted, fibroblasts can be isolated from relatively small skin biopsies, such as ear clippings (Seluanov et al., 2010), with minimally invasive sampling and harm to the individual. These cells can be expanded in tissue culture and, when immortalized, provide a highly scalable source of fresh DNA and RNA. Furthermore, living cells can be stored as frozen stocks in biobanks for decades (Fazekas et al., 2017). Tissue sampling can be done directly into the growth medium in ambient temperature, allowing several days for the transport to the laboratory without a need for dry ice or N_2_-cryocontainers, which greatly simplifies collection and logistics. The same applies to the isolated cells. As an example, we sent living cell suspensions from Joensuu, Finland to the ZFMK biobank in Bonn, Germany, over regular airmail in ambient temperature. The receiving laboratory can then amplify the cells by culturing and cryopreserve them in sufficient replicate stocks. It is worth to point out that the obtained genome assembly scaffolded to 23 autosomes, X and Y chromosomes, as expected for the species, shows no evidence of karyotypic drift caused by cell culture, dispelling possible concerns for the use of cell lines for such work (Gaertner et al., 2022).

The brown hare is an iconic European mammal, locally familiar to a wider public of nature-goers, farmers and hunters, while being at the same time an invasive species on many other continents. It has populations with complex ancestral makeup due to isolated Pleistocene refugia, frequent introductions and cross-species hybridization, making it an exceptionally interesting mammal for genetic studies. Furthermore, depending on the local environmental context, brown hares can be both an endangered as well as a rapidly expanding species, prompting opportunities for both conservation as well as invasion genetics studies. A high-quality reference genome assembly will allow us also to peer deeper into genome structure and chromosome evolution, which will be particularly interesting from the viewpoint of genetic introgression (Juric et al., 2016) and maintenance of species boundaries. Any issues of genetic compatibility between species should leave their fingerprint in the genomes (Skov et al., 2023), providing interesting insight into the speciation mechanisms (Wolf et al., 2010). Having a high-quality genome for both species has important implications for the fields of evolutionary and conservation biology. Due to the high contiguity of the genome assembly, it will be possible to perform accurate analyses of linkage disequilibrium as well as the identification of runs of homozygosity (ROH). Accurate ROH analysis is useful for identifying and understanding the impact of inbreeding burden on the genome of increasingly threatened and isolated species, as quantitative traits can be associated with ROH burden (Ceballos et al., 2018). Furthermore, this information might provide important insights into the hybridization of the brown hare with the mountain hare, as well as help to pinpoint genomic regions that might be helping the species to adapt to local environmental conditions during the ongoing range expansion (Pohjoismäki et al., 2021). Other *Lepus* reference genome assemblies available soon will help to complete this picture. Together, this genome assembly and the mountain hare genome assembly will provide a solid base for future studies.

## Acknowledgements

We acknowledge the DNA Sequencing and Genomics Laboratory, Institute of Biotechnology, University of Helsinki for the PacBio and Hi-C sequencing, with special thanks to Dr Petri Auvinen and Dr Martyn James for helpful discussions and personal interest towards the project. Dr Seppo Helisalmi and Dr Joose Raivo from the Gene diagnostics lab at Genome Center of Eastern Finland are thanked for providing the mtDNA sequencing service. We are grateful to the European Reference Genome Atlas (ERGA) for the opportunity to include our mountain hare genome into their pilot project. Dr Ann McCartney is thanked for her kind help throughout the process and Mr Jukka Pusa for his contribution in obtaining the brown hare specimen. We thank Ms Anita Kervinen for her valuable laboratory assistance and Dr. Mervi Kunnasranta for kindly allowing to use her hare photo in the publication.

## Data, scripts, code, and supplementary information availability

The genome assembly is available from the NCBI database under BioProject ID PRJNA950335. The cell line is available through the corresponding author as well as from the ZFMK biobank.

## Conflict of interest disclosure

The authors declare that they comply with the PCI rule of having no financial conflicts of interest in relation to the content of the article.

## Funding

This study belongs to the xHARES consortium funded by the R ‘Life initiative of the Academy of Finland, grant no. 329264.

## Notes

https://www.ncbi.nlm.nih.gov/biosample/SAMN33984520

### Competing Interest Statement

The authors have declared no competing interest.

### Summary of Updates

Included the PCI Genomics badge with embedded link to the recommendation. Updated annotation results and provided the accession number. Minor typos corrected and two references added regarding the history of the brown hare in Finland.

